# Transcending Markov: Non-Markovian Rate Processes of Thermosensitive TRP Ion Channels

**DOI:** 10.1101/2023.03.10.532104

**Authors:** Yuval Ben-Abu, Stephen J Tucker, Sonia Contera

## Abstract

The Markov state model (MSM) is a popular theoretical tool for describing the hierarchy of time scales involved in the function of many proteins especially ion channel gating. A MSM is a particular case of the general non-Markovian model, where the rate of transition from one state to another does not depend on the history of state occupancy within the system, i.e., it only includes reversible, non-dissipative processes. However, this requires knowledge of the precise conformational state of the protein and is not predictive when those details are not known. In the case of ion channels, this simple description fails in real (non-equilibrium) situations, for example when local temperature changes, or when energy losses occur during channel gating. Here, we show it is possible to use non-Markovian equations (i.e. offer a general description that includes the MSM as a particular case) to develop a relatively simple analytical model that describes the non-equilibrium behavior of the temperature-sensitive TRP ion channels, TRPV1 and TRPM8. This model accurately predicts asymmetrical opening and closing rates, infinite processes, and the creation of new states, as well as the effect of temperature changes throughout the process. This approach therefore overcomes the limitations of the MSM and allows us to go beyond a mere phenomenological description of the dynamics of ion channel gating towards a better understanding of the physics underlying these processes.

**Significance Statement:** Modeling ion channel processes has long relied on the Markovian assumption. However, Markov theory cannot translate situations in which the physical state of an ion channel changes during its gating process. By using a non-Markovian approach, we develop a simple analytical model that describes the non-equilibrium behavior of two temperature-sensitive TRP channels, TRPV1 and TRPM8. This model accurately describes and predicts their biophysical behavior as well as their temperature dependence. This approach therefore provides a better understanding of the physics underlying dynamic conformational changes such as those that occur during ion channel gating.

## Introduction

Proteins are the nanoscale orchestrators of life, performing diverse functions in living organisms. An example is their ability to regulate the selective movement of ions such as Na^+^ and K^+^ across biological membranes as part of the complex processes that control all forms of cellular electrical activity, especially the function of the heart and nervous system. A critical step in this process is the ability of specific protein pores or ion channels to change shape between their conductive and non-conductive forms. This ‘gating’ process allows them to regulate ion movement and couple it to a wide range of cellular signaling pathways [1].

Markov state models (MSMs) and their application to the conformational dynamics of biological systems [2] provides a theoretical approach that can describe kinetically relevant states and the rates of interconversion between them. Using this approach, many different types of protein conformational changes can therefore be examined [3] [4]. By stitching together a set of individual, short molecular simulation trajectories, MSMs therefore provide a summarized view of the ensemble of spontaneous fluctuations exhibited by proteins at equilibrium [5] [6], meaning that certain molecular information can be lost. A MSM is a particular case of the more general non-Markovian model. The term ‘non-Markov’ process or model covers all stochastic (random) processes with the exception of the small minority that can be described using the “Markov property”. In a MSM, the transition from one state to another depends only on the relevant transition rate from one state to the next, and does not include any history of state occupancy within the system, i.e. it only includes reversible, non-dissipative processes. The Markovian restriction is therefore useful because it facilitates a mathematical description of the process, but does not always fully describe the process being studied [7]. A particularly important limitation is that a MSM cannot describe the effect of temperature on a process [8]. In humans, the detection of both hot and cold temperatures by sensory neurons involves specialized temperature-sensitive ion channels such as members of the Transient Receptor Potential (TRP) family of channels [9]. The temperature sensitivity of these different channels spans the entire environmental range from noxious cold (<15°C) to injurious heat (>42°C) [10] [11] (**Figure 1A**) and allows the body to respond to a wide range of environments. One such example is TRPV1, which, as well as being activated by acidic pH and several chemical ligands, is also activated by supraphysiological temperatures (>42°C). Interestingly, capsaicin, one of the active ingredients in chilli peppers, is a natural agonist of TRPV1 and one of the main reasons that foods containing chilli taste ‘hot’. Another example is TRPM8, which has an opposite temperature activation profile to the TRPV1 being activated by decreased temperatures and also by the natural agonist, menthol [9].

**Figure 1:**
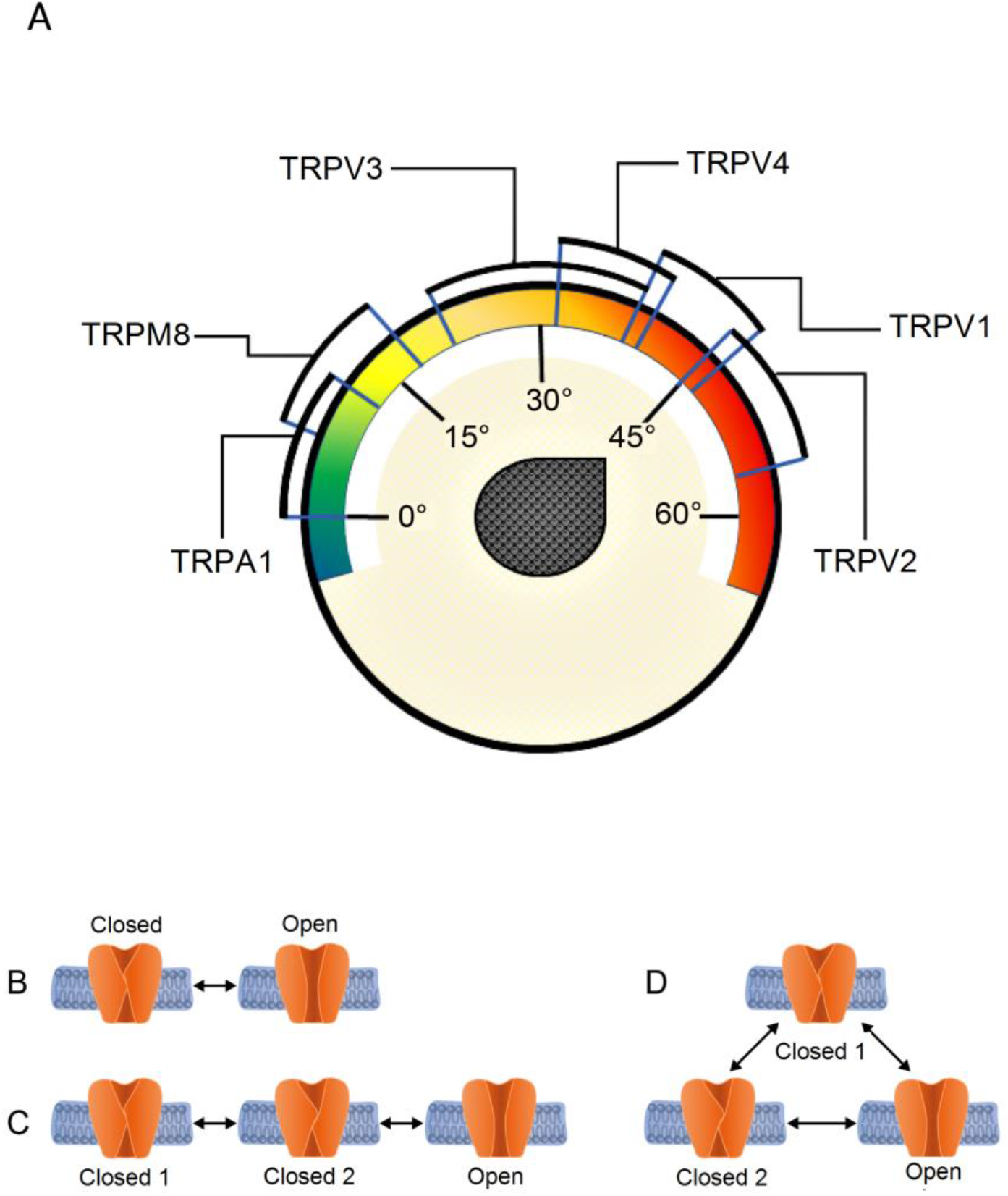
(A). Temperature sensitivity of different members of the Transient Receptor Potential (TRP) channel family. Schematic representations of two- (B) and three-state models (C, D) with time-dependent opening and closing rates. Each case represents a different situation predicted by a Markovian model.

The structure of many different thermosensitive TRPs have now been solved in a variety of conformational states [12] [13], but the molecular mechanisms that underlie their temperature-dependent regulation remain unresolved [12]. It has been proposed that gating-associated conformational changes result in changes in the heat capacity of the channel protein [14] and more recent studies have proposed a structural basis for the activation of TRPV1 by elevated temperature [15]. However, a mechanism that explains their activation by both hot and cold temperatures so far remains elusive. Undoubtedly, the rapid advances being made in CryoEM will provide important insights into the underlying structural basis of TRP channel gating, but the development of such hypotheses also relies upon the ability to compare such structural models of gating with accurate kinetic models derived from measurements of channel activity at different temperatures.

To address the limitations of the Markovian description, we propose a general non-Markovian model and demonstrate that this model can be solved analytically and used to model the behavior of TRPV1 and TRPM8 channels. This non-Markovian approach provides a theoretical framework for understanding the non-equilibrium thermodynamic activity of these channels, and highlights a new approach to understanding the physics of ion channel gating.

## Results

### Thermodynamic and kinetic limitations of the Markovian description

As with many types of proteins and ion channels, the kinetics of TRPV1 and TRPM8 gating are often described using MSMs [16] [17]. These Markov models are a simplified phenomenological representation of dynamic processes (e.g. channel gating) described quantitatively by a set of discrete states as they transition over time from one state to another. MSMs are common in the context of modeling ion channel activity as they switch between different conductive and non-conductive conformations such as those shown in **Figure 1B**. In such models, the complexity of real channel dynamics is reduced to a set of different kinetic states that represent discrete conformational states that a channel assumes at any given time. A number of rate constants are then used to define the equilibrium transitions between these different states [18].

However, in a MSM, the transition from one state to another does not take into account any history of state occupancy within the system, and so cannot predict realistic, nonequilibrium dynamics. MSMs also fail to account for irreversible processes such as temperature changes or energy dissipated during channel gating [7, 19, 20, 21]. More profoundly, the simplicity of MSMs limits the information that can be extracted and so they fail to be predictive unless the time-dependent conformational dynamics of the channel are known in detail. The principal shortcomings of a Markovian description of channel dynamics can therefore be summarized as follows:

i. MSMs require detailed knowledge of the structural changes that underlie the different kinetic relationships between states, and are traditionally expressed as arrows connecting these states (**Figure 1B-D**). As a result, MSMs do not make full use of experimental datasets because to use them, rather than comparing the data with the predictions of the model, the data must be forced on to the model [7] [22].
ii. In the case where there are fewer independent observations than parameters, MSMs are also not unique, meaning that many different models can fit the data [23] [24]. For example, as shown in **Figure 1D**, when there is more than one closed state in a system, six rate coefficients are required to define the model, but only four independent parameters are determinable from the data [7]. The same scenario also occurs in the two-state system (**Figure 1B**). The problem is usually solved by introducing symmetry within the system, i.e., certain rates are assumed to be identical and parallel or by obtaining orthogonal information via additional experimental approaches [7].
iii. MSMs assume that transitions occur from a finite number of states, whereas, in reality, there are likely to be several intermediate states [25], as well as fluctuating rates [26] and/or strong memory effects. However, when a system dissipates energy, it can never return to the exact same state as before because that state is now energetically different [27, 28]. For example, in the two-state model in **Figure 1B**, when the open channel returns to the closed state, since this process is energy-dependent, it can never assume the exact same closed state it was in before; it must therefore become a different (albeit similar) closed state [7] [20] [29].
iv. Finally, the classical Markovian equation describing channel kinetics normally assumes a constant temperature throughout the process, *i.e*. the channel is not energetically connected with its interfaces within the membrane and surrounding fluid. Reported attempts to model temperature changes in the system require the introduction of at least seven or eight additional states [7] [20] [30]. However, real channel proteins are highly dynamic with some parts undergoing large-amplitude Brownian motions and other rearrangements [31] [32] and hence the energy of these different states is not constant. Therefore, even when modified, Markov models have important practical and conceptual limitations [22] [33].

To address these limitations, we now introduce a general non-Markovian model that goes beyond the Markovian nonlinear stochastic systems described in **Figure 1** to include non-Markovian descriptions of Brownian motion [22] [20] [31] [32]. This non-Markovian model provides a theoretical framework for understanding the nonequilibrium thermodynamic activity of these channels [14] [17] [22], and other channels in general, without the unrealistic limitations of Markovian approaches. We further test this model on experimental data [34] [35] [36].

### Underlying theory for a non-Markovian description of channel dynamics

In the following section, we summarize our non-Markovian model. The whole mathematical description is given in the **Supplementary Information**. For clarity, we first consider a simple model with two states, which is then extended to three states case (**Figure 1**)

First, we present a solution for the infinite case, as is usually done in MSMs, and then use our model to solve the problem of temperature changes during channel activity. To find states that define the conformational changes we are interested in amongst the infinite sample space of all possible conformations, we use Borel Sigma Algebra. This is a tool from modern probability theory that is useful for mathematics with uncountable sample spaces. The most important result is that our mathematical derivation allows a direct comparison with experiment, by giving a set of equations that relate thermodynamic quantities obtained from experimental data to our non-Markovian model predictions.

First, our system is defined as a stochastic continuous system having the following special parameters: homogenous time t, continuous state times {*θ_i_*}, *a* certain probability space (*Ω* = *B, P*), expectation (E), occupation times for states 1 and 2, variation Var (Borelian probability groups) and discrete space of values (states); all these parameters are described in detail in the **Supplementary Material**. After defining this stochastic system, we defined and found the continuous distribution. Next, we defined a continuous random process *ξ* = *ξ(t)* as a state in the system at time *t, where t ≥ 0*. We assume that at *t=0 ξ(0) = 0*

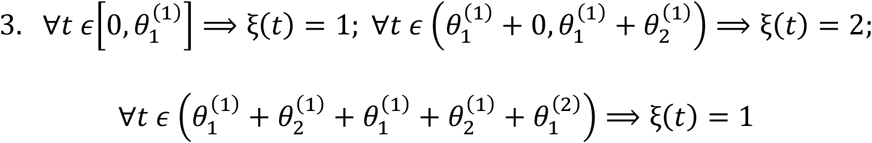

etc. Symbolically (the full description appears in the **supplementary data**):

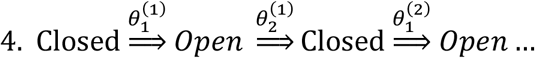

In the case of three states, m=3 (we call the states “Closed1”, “Closed2” and “Open”) and *ξ*(*t*) = 3, *t* ≥ 0. This case presents two possibilities symbolically in two states:

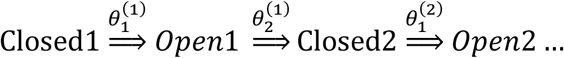

And in three states:

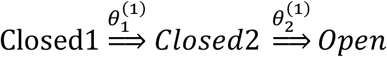

From these two and three states as elaborated in the **Supplementary Information**, we solved the setup model using the numerical method described by Goychuk et al. [7, 20]. Since these methods deal with a stochastic system, the involvement of stochastic resonance is necessary [20]. Therefore, for this, we analyzed numerical calculations of experimental data for TRPV1 and TRPM8 which are activated by heat and cold respectively. Unlike standard Markovian modeling, we assume that the transition from Closed1 to Closed2 is not rate-limited but is instead, characterized by a broad distribution of rates. This transition then continues on to a new state that implies the gating process soon becomes non-Markovian within two or three states.

If our model is to reproduce the experimental results it must demonstrate that the rates of transition from closed to open, and *vice versa*, are asymmetrical in the nonequilibrium infinite process, i.e. it must describe a realistic channel out of equilibrium that loses energy and whose energy losses affect the rate of the transitions and the kinetic reaction constant. Therefore, according to Goychuk et al. [7] [20], together with our theory, we applied this using these relationships for the kinetic reaction constant *k_o,c_*(*t*) (where *o*, means open and *c* means close):

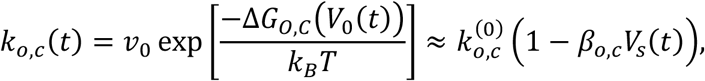

Where, in the absence of driving:

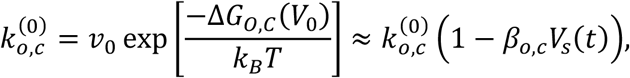

With Δ*G_O,C_*(*V*_0_) denoting the corresponding static-free energy barriers, *V_0_* being the static voltage in the absence of a signal, i.e., *V*(*t*) = *V*_0_ + *V_s_*(*t*) and 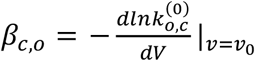, *k_B_* is the Boltzmann constant and T is the absolute temperature. We continued to develop our theory (see **Supplementary Information**) and arrived at the following expression and application of ∈(T) by description of the temperature sensitivity, free energy, rates and the other thermodynamics parameters in:

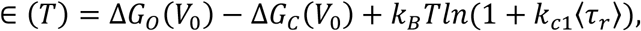

Whereby Δ*G_O_*(*V*_0_) = *G*^#^ – *G_c_* and Δ*G_O_*(*V*_0_) = *G*^#^ – *G*_o_, where *G*^#^is the free energy of the transition state and Δ*G_O_* = *H_c,o_* – *TS_c,o_* is the free energy of the open (closed) conformation, and then:

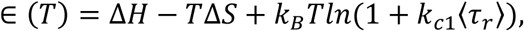

where Δ*H* = *H_O_ – H_C_* is the difference between the thermodynamic enthalpies of the open and closed conformations and Δ*S* = *S_O_* – *S_C_* is the corresponding entropy difference. Importantly, the results match the typical experimentally-determined responses for these channels (**Figure 2**) along with the non-Markovian behavior as described in the **Supplementary Information** and **Figure 3**.

**Figure 2:**
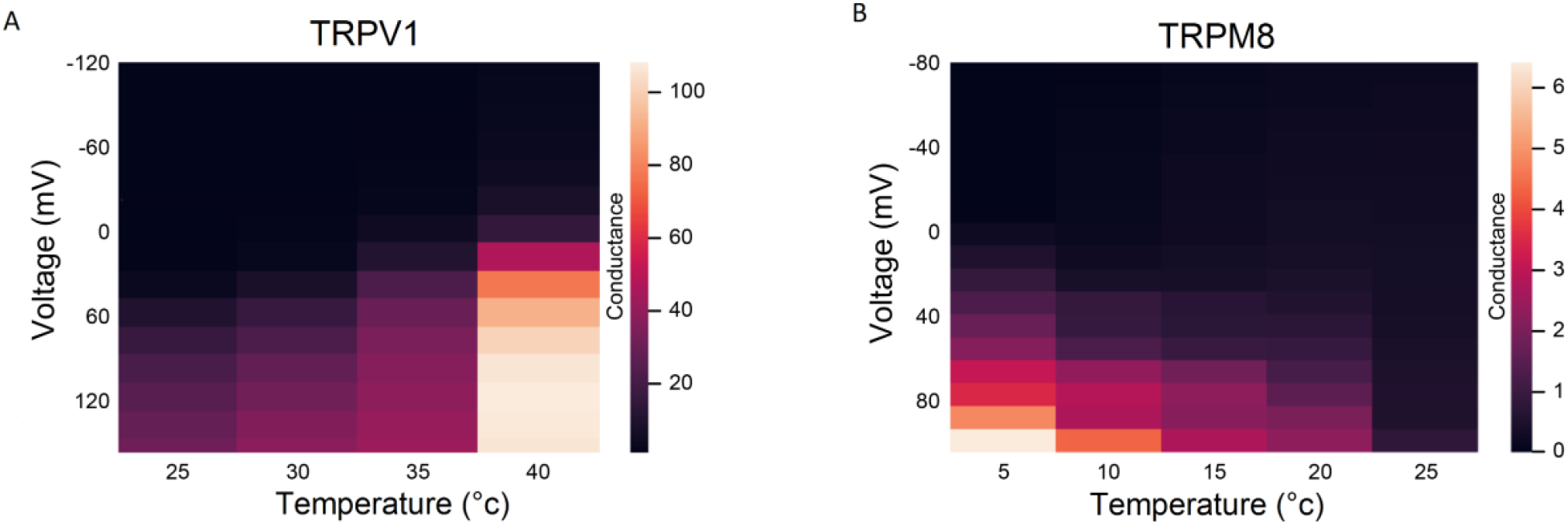
Typical temperature sensitive behavior of the TRPV1 (A) and TRPM8 (B) channels as determined by experimental measurements of channel activity at different voltages. Conductance-voltage (G-V) relationship of the plateau current for the wildtype TRPV1 and TRPM8 channels are shown at different temperatures. This shows the typical increase in TRPV1 channel activity associated with increased temperatures, and vice-versa for TRPM8.

**Figure 3:**
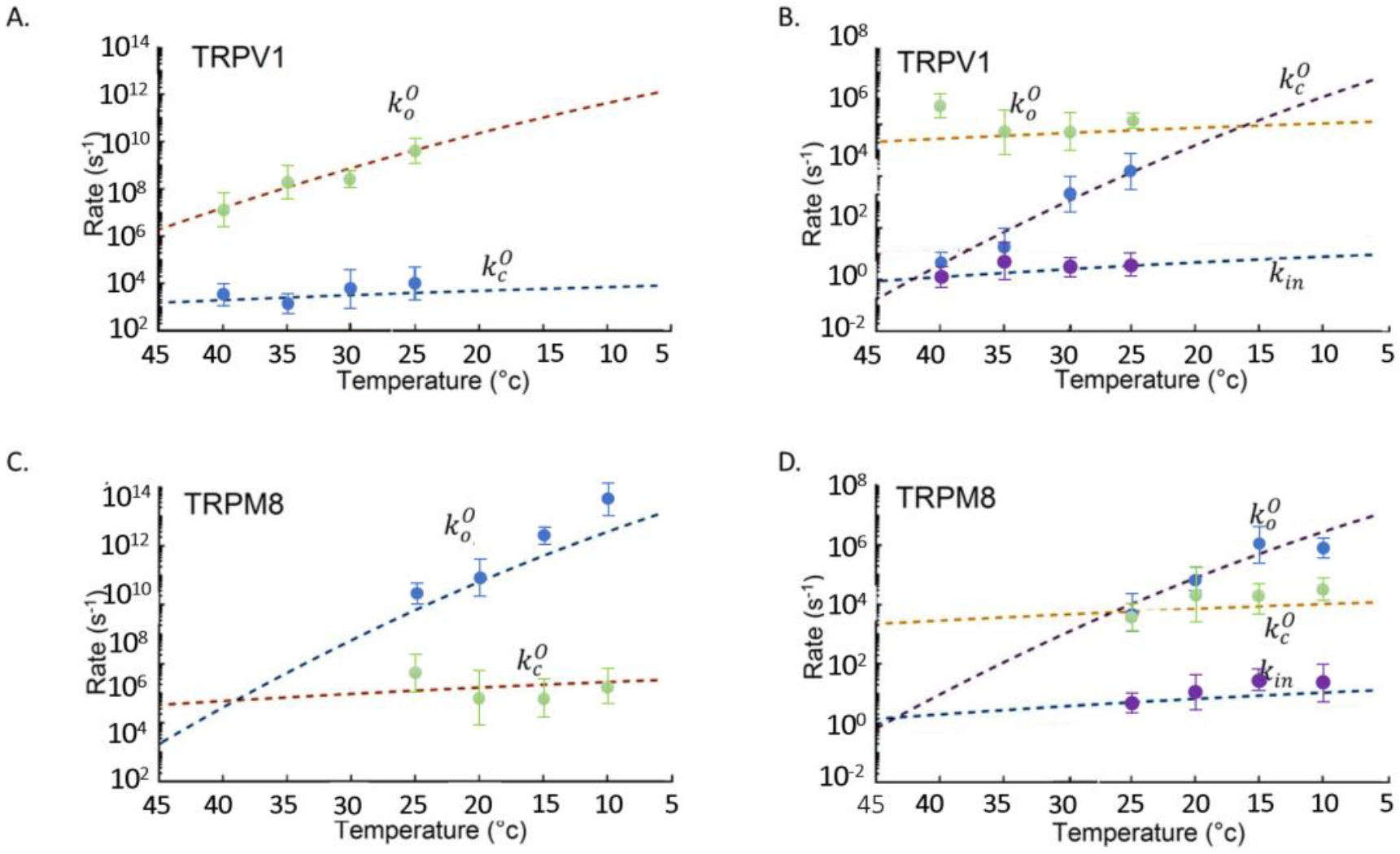
The experimentally determined temperature-dependent responses shown in Figure 2 were analyzed and modeled based on the equations appearing in the results section and in the Supplementary Materials and according to Guychuk et al. [29]. A and C represent the two-state model whereas B and D represent the three-state model. The dots in different colors represent the opening and closing constants of TRPM8 and TRPV1 channels. These points were found to be adapted to the non-Markovian model shown in the dashed line, according to two main states in each of the two-state and three-state channels. The corresponding two-state non-Markovian opening rate is defined as 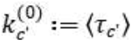, where 〈*τ’*〉 is the mean residence time. The rate 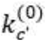 denotes the transition from the open state O to the closed state C.

After a series of numerical calculations based on the theory appearing in the **Supplementary Information**, we next calculated the temperature-dependence of the corresponding transitions of both channels with either two or three states. **Figure 3** presents this two- and three-state non-Markovian temperature-dependent analysis of TRPV1 (A and B) and TRMP8 (C and D). Interestingly, the corresponding two-state non-Markovian opening rate is defined as 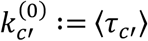, where 〈*τ_C’_*〉 is the mean residence time. The rate 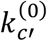 denotes the transition from the open state O towards the closed state C.

### Application of a non-Markovian model to TRPV1 and TRPM8

Using the mathematical non-Markov model described above, we performed numerical calculations for the experimental dataset obtained from TPRV1 and TPRM8 channels. These descriptions were chosen to mimic the experimental temperature dependence of the TPRM8 and TPRV1 heat- and cold-sensitive channels. For both channels, these datasets relate to activity recorded from heterologously expressed channels over the range of voltages and temperatures shown in **Figure 2** [34, 35, 36]. Thermodynamic parameters such as enthalpy (ΔH) and entropy (ΔS) were also derived from this data and analyzed within a framework of two- and three-state non-Markovian dynamics based on Markovian equations. This non-Markovian methodology is summarized below and further developed in the **Supplementary Information**.

### Replication of thermosensitivity by this non-Markovian model

Using our approach, we calculated two models for TRPV1 and TRPM8 gating based on the two- and three-state models described above (*Closed ↔ Open* and *Closed1 ↔ Closed2 ↔ Open*). **Figure 3** and **Figure 4** represent the data obtained from applying our non-Markovian approach to the simplest two-state model (*Closed ↔ Open*) that we present in the second part of the **Supplementary Material**. The results show that the rate coefficients *k_o_* and *k_c_* are non-symmetrical (unlike the MSM prediction) and that the temperature sensitivity of the channel can be correctly replicated.

**Figure 4:**
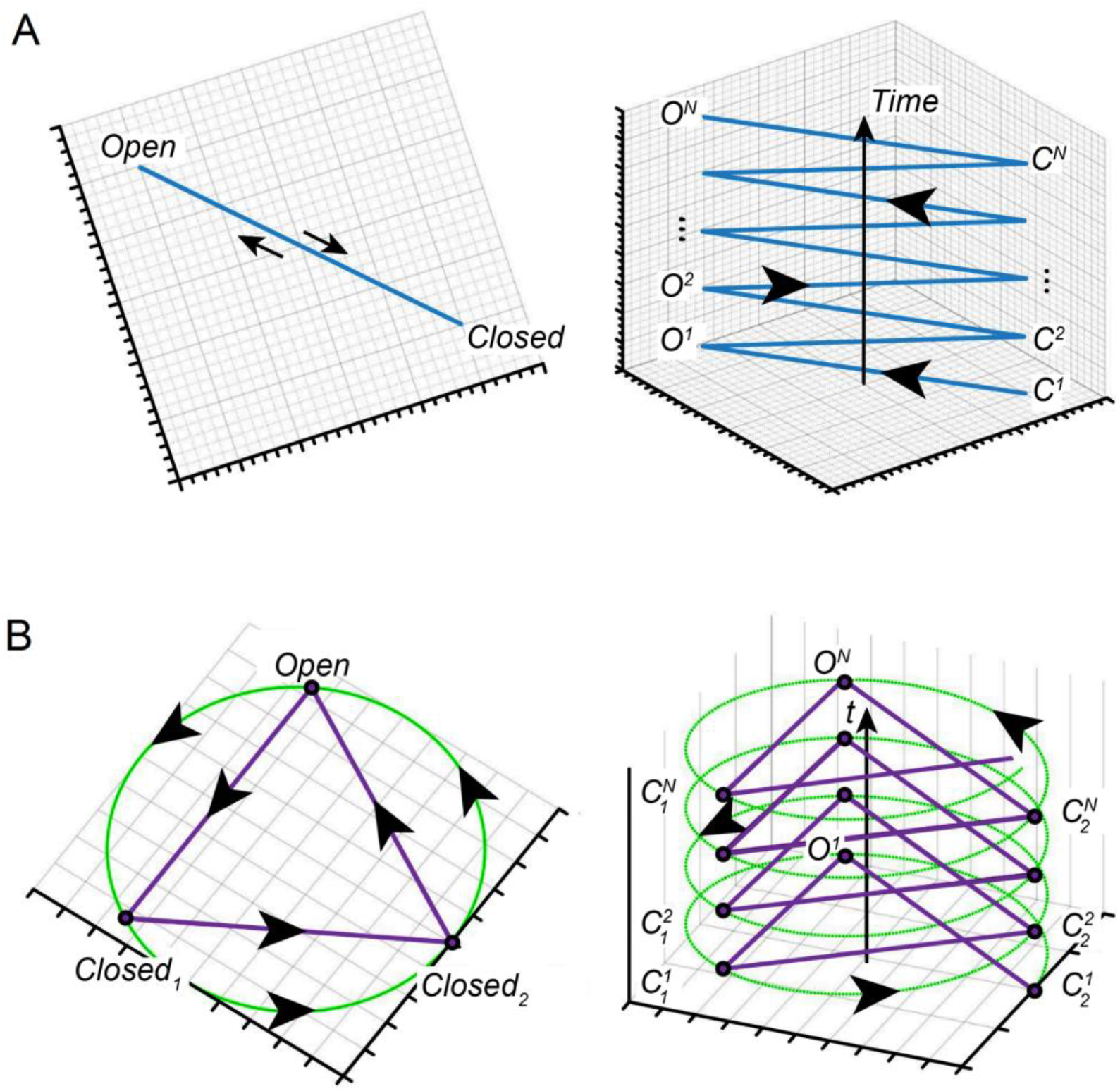
Cyclical graphs of (A) the two model and (B) the three state models. This type of system describes the trajectory of each case, which is given by a random arrangement of open and closed states over time. Both descriptions allow visualization of the infinite cases represented by our model. The left panel in each model state represents the top view and the right panel the side view. This model represents the infinite process whereby each state is new and is not similar to the last one. This is different to what would be represented in a Markovian model.

Next, we analyzed the three-state model (*Closed1↔ Closed2 ↔ Open*) using our non-Markovian model, then transformed it into a practical model. This practical model is based on Δ*G*, Δ*S and* Δ*H* calculations from the dataset of the three states, *Closed1, Closed2* or *Open* of TRPV1 and TRPM8 channels. The results of this analysis are shown in **Figure 3 B, D** and **Figure 4**

Importantly, our model correctly reproduces the asymmetry in the rate coefficients *k_o_* and *k_c_*, as well as the thermosensitivity of both channels i.e. it reproduces TRP channel activation by both heat and cold. Indeed, in both cases shown in **Figure 3**, the non-Markovian models can clearly predict the states of the channel with greater accuracy than can be done using a classical Markov model and can accommodate a wide range of temperature changes, asymmetrical rate coefficients and the generation of rate constants over a potentially infinite period of time.

## Discussion

In this study, we have shown that it is possible to use non-Markovian equations to develop a relatively simple analytical model that describes the non-equilibrium behavior of the thermosensitive TRP channels, TRPV1 and TRPM8. This model accurately predicts many key aspects of their behavior, in particular the effect of temperature changes on channel activity. This approach therefore overcomes the limitations of traditional Markovian models and allows us to go beyond a mere phenomenological description of the dynamics of ion channel gating.

MSMs have been particularly useful to interpret the dynamics of many K^+^ and Na^+^ channels [37] [38] but have particular limitations when it comes to modelling the effects of temperature [7]. For example, the gating dynamics of the voltage-gated *Shaker* K^+^ channel have been reasonably described at 20°C by an eight-state Markovian scheme including seven closed state [7, 37]. However, to accommodate the effect of even a small range of temperatures (10-20°C) requires three additional closed states [39]. The proliferation of discrete states, albeit necessary for the description of experimental data, is rarely addressed within Markovian descriptions [20]. To explore these issues, we therefore examined thermosensitive gating using the non-Markov model described in the **Supplementary Material**.

One of the specific issues that it addresses is that of energy dissipation during gating. For example, looking at a simple process such as *A ↔ B*, a Markov model predicts that *A* becomes *B* and then returns to the exact same state, *i.e. A* [40]. However, in reality, *A* becomes *B*, but *B* cannot return to exactly state *A* again because the overall process loses energy, and so when *B* returns to state *A* it is not exactly the same as it was before. This means that some new variant of state *A* is produced, and although its functional properties may be similar, the physicochemical nature of the state is formally different. [40] [41] For that reason, a description of such “different” or “new” states with MSMs requires a more complex calculation for each state and conceptualizes the overall process as being finite. However, this non-Markovian approach is more accurate as it can predict the dynamics of an essentially infinite process.

Simple MSMs such as *Closed ↔ Open* can be refined by creating additional states, but the number that can be realistically added are limited. However, the non-Markovian approach provides a potential solution to this and works as follows:

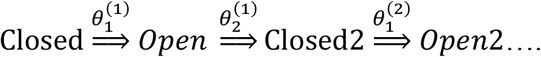

Importantly, this non-Markovian model is an infinite process whereby each open or closed state may be similar to the previous version, but formally each state is defined as new. This situation is therefore closer to what occurs in reality since energy is lost in every transition.

Of course, Markov models are still informative, but are theoretically limited. The same limitation also occurs in the three-state MSM described in **Figure 1** and **Figure 3B, D** where there is an additional closed state. In this example, a new manipulation was performed to solve the case and the process is described as:

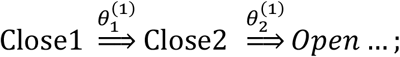

For this equation, we analyzed experimental data for TRPV1 and TRPM8 and got complete agreement to the model (**Figure 3**), i.e. the channel transitions were perfect according to non-Markovian behavior and we can see the accuracy of the constant reaction rate from one state to another. By contrast, a traditional Markov model of such a process involves describing transitions between the “same” states without any energy loss or any other thermodynamic parameters. However, the effects of temperature on channel activity is an ongoing process that depends on many factors such as reaction constants, entropy and enthalpy. As we can see in **Figures 3A and 3B**, this non-Markovian approach is accurate in predicting the temperature-dependence of the constant reaction rate from one state to another, together with the values of the rate constants, and all the new states that occur in the process. Furthermore, this is done without adding any additional states to the equations as would likely be required for a MSM of this process [7] [20] [31].

We therefore present a new analytical approach for describing the complex gating dynamics of TRPV1 and TRPM8 channels based on a new general non-Markovian model that overcomes the important limitations of MSMs. Despite the relative complexity of our non-Markovian approach, it is still relatively easy to implement in practical examples, as we have described here for TRPV1 and TRPM8. Our model is able to accurately reproduce experimental results, including the fact that the rates of transition from closed to open, and vice versa, are asymmetrical in the non-equilibrium infinite process. Our approach is therefore able to describe a realistic channel out of equilibrium that loses energy and whose energy losses affect the rate of the transitions and the kinetic reaction constant. Finally, this model reproduces the effects of temperature on the gating process and is therefore particularly relevant for thermosensitive channels such as TRPs where an accurate description of thermodynamic processes is critical. Moreover, from a purely practical point of view, the simple two and three state systems shown in **Figure 1** can be expressed as a finite collection of states in which each open or closed state can be significantly different to their previous versions as time evolves. The system can therefore be described as an evolution graph in time, interpolating between the different states of the system. An organized graph, such as the ones presented in **Figure 4**, for these two examples and may be given by an undirected state graph, where the evolution trajectory is given by a random arrangement of open and closed states over time. A generalized model is also presented in **Figure 5** that compares between two and three states model description of Markov and Non-Markovian model. By this model, we can see the infinite process that each state is new and is not similar to the last one as represented in the Markov model.

**Figure 5:**
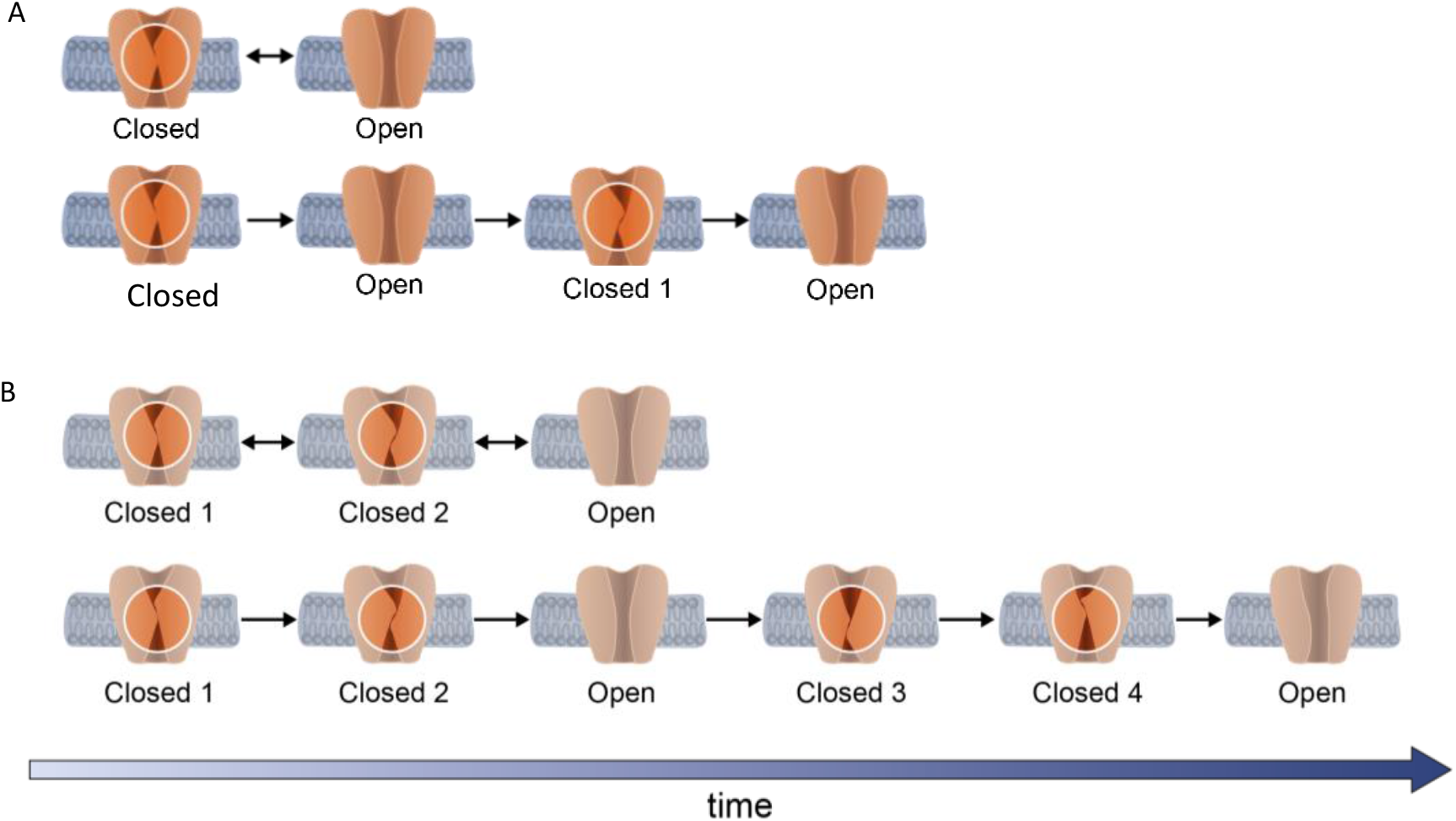
Comparison between the two- (A) and three-state (B) descriptions of the Markovian (first part of A and B) and the non-Markovian models (lower part of A and B). This model describes the infinite process whereby each state is new and is not similar to the last one, as is represented in the Markovian model in the first panel in A and B.

In conclusion, this model accurately describes and predicts many key aspects of TRP channel behavior, and overcomes the limitations of traditional Markovian models thereby allowing us to go beyond a mere phenomenological description of their gating kinetics. By adapting this methodology, we expect this approach can be used to help understand the behavior of many different ion channels as well as many other types of dynamic proteins.

## Materials and Methods

### Experimental datasets

the electrophysiological data for TRPV1 was generously contributed by Prof Avi Priel (Hebrew University of Jerusalem, Israel) and for TRPM8 by Prof Jorg Grandl (Duke University, USA). These standard datasets of channel thermosensitivity were obtained as previously described for these channels [36] [42].

### Data analysis and statistical tests

Analysis was performed with Excel (Microsoft, USA). Graphs were made using Prism (GraphPad Software, USA), Matlab (MathWorks, USA) and Mathematica (Wolfram Research, USA) software. Electrophysiological datasets were analyzed with Igor Pro (WaveMetrics, USA).

## Notes

**Competing Interest Statement:** The authors declare no competing interests.

### Competing Interest Statement

The authors have declared no competing interest.

