## Supplementary Material for "Transcending Markov: Non-Markovian Rate Processes of Thermosensitive TRP Ion Channels"

### Models for 2 and 3 states: setup

Here, we develop a non-Markov model that can be used to interpret the behavior of ion channels. First, we will consider a model for 2 states, and then we will extend it to 3 states.

Individual states are divided into groups of "open" and "closed" states; the channel conducts ions only in an "open" state. The two-state model consists of one closed (C) and one open (O) state, while the three-state model consists of $C_1$ (Closed1) $C_2$ (Closed2) and O (Open). The description of ion channel transitions in the Markov model has many limitations [10] [28]. Here, according to the theoretical model we transformed into practical model according to $\Delta G, \Delta S$ $and$ $\Delta H$ dataset calculation of the three states Closed1, Closed2 or Open of TRPV1 and TRPM8 channels.

### Model theory:

### Two state case

In this section, we consider the case where m = 2, i.e., M = M(2) = {1, 2. }; $\theta_1$ and $\theta_2$ are the occupation times for states 1 and 2, respectively. More precisely, the occupation times of state 1 are the sequences of the independent positive random variables - s $\{\theta_1^{(j)}\}$ having at the same continuous distribution as $\theta_1$:

$$\mu(A) = P(\theta_1^j)\epsilon A) = P(\theta_1 \,\epsilon\, A)$$

Analogously, $\theta_2$ is the occupation time of state 2; the occupation times of state 2 are the sequences of independent positive random variables - s $\{\theta_2^{(j)}\}$ having at the same continuous distribution as $\theta_2$ :

$$v(B) = P(\theta_2^j)\epsilon B) = P(\theta_2 \,\epsilon\, B)$$

Here, *A* and *B* are arbitrary Borel sets on the right-hand side of the real axis. Let us define now – a continuous random process $\xi = \xi(t)$ as a state of described above system at time $t;\ t \geq 0$. In detail, one can suppose that $\xi(0) = 0$, and

$$\forall t \,\epsilon \left[0, \theta_1^{(1)}\right] \Rightarrow \xi(t) = 1;\ \forall t \,\epsilon \left(\theta_1^{(1)} + 0, \theta_1^{(1)} + \theta_2^{(1)}\right) \Rightarrow \xi(t) = 2;$$

$$\forall t \,\epsilon \left(\theta_1^{(1)} + \theta_2^{(1)} + \theta_1^{(1)} + \theta_2^{(1)} + \theta_1^{(2)}\right) \Rightarrow \xi(t) = 1$$

etc. Symbolically

$$\text{Closed} \overset{\theta_1^{(1)}}{\Longrightarrow} Open \overset{\theta_2^{(1)}}{\Longrightarrow} \text{Closed} \overset{\theta_1^{(2)}}{\Longrightarrow} Open \ldots$$

Therefore, let us define the probabilities distribution of $\xi(t)$:

$$p_1(t) := P\xi(t) = 1;\ p_2(t) := P\xi(t) = 2) = 1 - p1(t)$$

So that:

$$p_1(0) = p_1(0+) = 1$$

Introduce also a regeneration moment:

$$\tau := \theta_1^{(1)} + \theta_2^{(1)}{}_1$$

The distribution $\zeta$ of the random variables $\tau$ is the convolution of ones for $\theta_1^{(1)} and\ \theta_2^{(1)}$:

$$\zeta(C) = \mu * v\ (C), C \subset R_+^1$$

$$\zeta(C) = \mu * v\ (C) = \int_{R^2} I_C\ (x + y)\mu(dx)v(dy)$$

and $I_c(\cdot)$ is the indicator function of the measurable set C. Recall that the distribution of sums of two random independent variables is equal to the convolutions of summands. Of course, the one-dimensional distributions may be described by their distribution functions, i.e.

$$F(t) = P\ (\theta_1 < t), G(t) = P\ (\theta_2 < t), t \geq 0$$

We use the notation:

$$\bar{F}(t) = 1 - F(t) = P\ (\theta > t), t > 0;$$

The addition function of distribution, which is used often in the reliability theory, mean the connection define here as:

$$H(t) = F * G(t) = \int_0^t F(t-y)G(dy) = P\ (\theta_1 + \theta_2 < t)$$

The convolution between the function $v = v(t)$ and measurement of $v$ is defined as

$$v * v(t) = v * t(t) \stackrel{\text{def}}{=} \int_0^t v\,(t-s)v(ds)$$

Using the renewal **theorem 2.1** of the Volterra Integral equation [1-5], the function $p_1 = p_1(t)$ is a unique solution in the space of all bounded measurable functions that follow the ordinary norm $\|p(\cdot)\|\infty \stackrel{\text{def}}{=} \sup_{t\geq 0}|p(t)|$ of the following Volterra's integral equation:

$$p_1(t) = 1 - F_{\theta_1}(t) + \int_0^t p_1\,(t-s)F_{\theta_1+\theta_2}(ds)\ = 1 - F_{\theta_1}(t) + \int_0^t p_1\,(t-s)H(ds)$$

Or equivalently

$$p_1(t) = 1 - F_{\theta_1}(t) + [p_1 * H](t)$$

The proof is based on the classical method of moment regeneration: it can be calculated using the formula:

$$\tau = \tau_{reg}\theta_1^1 + \theta_2^1 = \theta_1 + \theta_2$$

Namely, we find it by means of the full probability formula

$$P(\xi(t) = 1) = \int\int_{R_+^2}^t P\left(\xi(t) = \frac{1}{\theta_1} = x, \theta_2 = y\right) F_{\theta_1+\theta_2}(dx, dy) = I_1 + I_2$$

where the partition of the integral has the form:

$$I_1 = \int\int_{x,y\geq 0, x+y>t}, I_2 = \int\int_{x,y\geq 0, x+y\leq t}$$

And we have:

$$I_1 = \int\int_{x,y\geq 0, x+y>t} P\left(\xi(t) = \frac{1}{\theta_1} = x, \theta_2, = y\right) F_{\theta_1+\theta_2}(dx, dy) = P(t < \theta_1) = 1 - F_{\theta_1}(t)$$

Further:

$$I_2 = \int\int_{x,y\geq 0, x+y\geq t} I(t > x, t > x+y) F_{\theta_1+\theta_2}(dx, dy) =$$

$$\int\int_{x,y\geq 0,x+y\geq t} I(t > x + y)F_{\theta_1+\theta_2}(dx, dy) =$$

$$\int\int_{R_+^2} I(t - x - y)I(x + y \leq t)F_{\theta_1+\theta_2}(dx, dy) =$$

$$E_{P1}(t - \theta_1 - \theta_2)I(\theta_1 + \theta_2 \leq t) =$$

$$I_2 = \int_0^t p_1(t - s)F_{\theta_1+\theta_2}(ds) = \int_0^t p_1(t - s)H(ds) = [p1 * H](t)$$

Therefore, to investigate the asymptotical behavior as $t \to \infty$ of the solution $p_1(t)$ of the equation:

$$\forall t \in \left(\theta_1^{(1)} + \theta_2^{(1)} + \theta_1^{(1)} + \theta_2^{(1)} + \theta_1^{(2)}\right) \Longrightarrow \xi(t) = 1$$

we use the Tauberian theorem (a theorem that deduces the convergence of a series on the basis of the properties of the function it defines and any kind of auxiliary hypothesis which prevents the general term of the series from converging to zero too slowly),using the Laplace transform $\Lambda$ , which is defined for any (measurable) bound function $f(t), t \geq 0$

$$\Lambda[f](\lambda) \stackrel{\text{def}}{=} \int_0^\infty e^{-\lambda t} f(t)dt, \Re\lambda > 0$$

Here and in the sequel, $\Re\, z$ denotes an ordinary the real part of the complex number z, so we allow the number $\lambda$ to be complex. The Laplace transform for the arbitrary non-negative distribution, μ, is defined as follows:

$$\Lambda[\mu](\lambda) \stackrel{\text{def}}{=} \int_0^\infty e^{-\lambda t}\, \mu(dt)$$

So that if the measure μ is distribution of the non-negative r.v. $\beta$, i.e. $\mu(A) = P(\beta \epsilon A)$, then evidently $\Lambda[\mu](\lambda) = Ee^{-\lambda\beta}$. This integral converges for the complex values $\lambda$ for which $\Re\, \lambda \geq 0$.

Of course, if the distribution $\mu$ has density dμ/dt = y(t), then both definitions coincide.

We recall the following important property: the Laplace transform is closely and continuity related to the source distribution to a weak convergence and we know that

$$\Lambda[\mu * v](\lambda) = \Lambda[\mu](\lambda) \cdot \Lambda[v] \cdot (\lambda)$$

In the **theorem 2.2**, suppose $0 < E\theta_1 < \infty,\ \ 0\ < E\theta_2 < \infty.$ Then

$$\lim_{x\to\infty} p1(t) = \frac{E\theta_1}{E\theta_1 + E\theta_2}$$

The proof is that one can assume without loss of generality that there exist a probability density function $h(t) = dH(t)dt.$ We apply the Laplace transform on both sides of the equation as mentioned above:

$$p_1(t) = 1 - F_{\theta_1}(t) + \int_0^t p_1\,(t-s)F_{\theta_1+\theta_2}(ds)\ = 1 - F_{\theta_1}(t) + \int_0^t p_1\,(t-s)H(ds)$$

And we get

$$\Lambda[p1](\lambda) = \Lambda[1 - F_\theta](\lambda) + \Lambda[p1](\lambda)\cdot\Lambda[h](\lambda),$$

Following

$$\Lambda[p1](\lambda) = \frac{\Lambda[1 - F_\theta](\lambda)}{1 - \Lambda[h](\lambda)}$$

We deduce consequently that as $\Re\,\lambda \to 0+$

$$\Lambda\big[[1 - F_\theta](\lambda)\big]\sim\int_0^\infty [1 - F_{\theta 1}](t)dt = E\theta 1^{-\lambda t}$$

$$1 - \Lambda\big[[h](\lambda)\big]\sim\lambda\cdot[E\theta 1 + E\theta 2]$$

$$\Lambda[p1(\lambda)]\sim\frac{E\theta 1}{\lambda[E\theta 1 + E\theta 2]}$$

**2.3 The three-state case**

Here, we derive the model for a more complex case with m=3, i.e., the system has 3 states, $\xi(t) = 3, t \geq 0.$ This case presents 2 possibilities, symbolically:

$$\text{Closed1} \overset{\theta_1^{(1)}}{\Longrightarrow} Closed2 \overset{\theta_2^{(1)}}{\Longrightarrow} Open\ldots\ldots$$

$$\text{Closde1} \overset{\theta_1^{(1)}}{\Longrightarrow} Open1 \overset{\theta_2^{(1)}}{\Longrightarrow} \text{Closed2} \overset{\theta_1^{(2)}}{\Longrightarrow} Open2 \ldots$$

The first has a probability, $q_+ = q_r$; the second has a probability $q_- = q_1$; $0 < q_- < 1$; $q_+ + q_- = 1.$ Independently, on the random variables – s $\{\theta_j^k\}$ which are also independent. If we consider that the random variables $\xi(t)$ are homogenous and Markovian, then automatically both the switch probabilities $q_-, q_1$ are proportional to the inverse exception of the values $\theta_1$ and $\theta_2$. We now offer (by analogy with the Markovian case) in the general case of non-Markovian process $\xi(t)$ the following switch probabilities:

$$q_1 := \frac{E\theta_2}{E\theta_1 + E\theta_2}, \qquad q_{r,0} := \frac{E\theta_1}{E\theta_1 + E\theta_2},$$

Let us define the following important probabilities:

$$p_1(t) := P(\xi(t) = 1);\ p_2(t) := P\xi(t) = 2) = p_3(t) := P(\xi(t) = 3)$$

Where, of course

$$p_1(t) + \ p_2(t) + p_3(t) = 1$$

The initial condition $p_1(0) = \ p_1(0+) = 1$. The corresponding occupation times will be denoted as before:

$$\theta_j = \left(\theta_j^{(1)} + \theta_j^{(2)} \ldots\ldots\ldots..+\theta_j^{(k)}\right) j = 1, k = 1,2,3, \ldots.$$

Let us introduce the random variable $\zeta$ as the first time after $\theta_1$ when the random process $\xi(t)$ returns to the initial state (state one):

$$\zeta = \ \zeta_1 = inf\left\{t, t > \theta_1^{(1)}, \xi(t) = 1\ \right\} - \theta_1^{(1)}$$

**Theorem 3.1.** The distribution

$$\mu_\zeta = q_- \cdot q_{\theta_2} + q_+ \cdot q_{\theta_2+\theta_3} *\ \mu_\zeta$$

**Proof.** Again, the relation follows immediately again from the full probability formula as long as, after the occupation time spent in the second state, random process $\xi(t)$ returns to the first state with probability $q_-$ and switches to the third state with additional probability $q_+ = 1 -$

$q_-$, whence it returns to the second state after time $\theta_3^{(1)}$ (recursion). As a corollary, we conclude applying the Laplace transform:

$$\Lambda[\mu_\zeta](\lambda) = \frac{q_- \, \Lambda[\mu\theta_2](\lambda)}{1 - \, q_+ \cdot \Lambda[\mu\theta_2](\lambda) \cdot \Lambda[\mu\theta_3](\lambda)}$$

After simple calculations

$$\mathrm{E}\zeta = \frac{E\theta_2}{q_-} + \frac{q_+ + E\theta_3}{q_-}$$

If of course, both expectations $E\theta_2$ and $E\theta_3$ exist (are finite).

**Theorem 3.2**. the function $p_1 = \ p_1(t)$ is the unique solution of the following Volterra integral equation of renewal type:

$$p_1(t) = 1 - F_{\theta_1}(t) + \int_0^t t_1(t-s)F\zeta(ds)$$

Proof is quite analogous to one in theorem 2.1; it is sufficient to note that here for the random process $\xi(t)$, the moment of regeneration is equal to $\zeta$. Therefore,

$$\Lambda(p_1)(\lambda) = \frac{\Lambda[1 - F_{\theta_1}](\lambda)}{1 - \, \Lambda[\mu_\zeta](\lambda)}, \Re\lambda > 0$$

If all three $E_{\theta_j}, j = 1, 2, 3$ are finite, then

$$\exists \lim_{t\to\infty} (p_2)(t) = \frac{E\theta_1}{E\theta_1 + E\zeta} =$$

$$\frac{E\theta_1}{E\theta_1 + \frac{E\theta_2}{q_-} + q_+ \frac{E\theta_3}{q_-}}$$

The other probabilities $(p_2)(t), (p_3)(t)$ may be found analogously:

$$\lim_{t\to\infty} (p_2)(t) = \frac{E\theta_2/q_-}{E\theta_1 + \frac{E\theta_2}{q_-} + q_+ \frac{E\theta_3}{q_-}}$$

$$\lim_{t\to\infty} (p_2)(t) = \frac{q_+ E\theta_2/q_-}{E\theta_1 + \frac{E\theta_2}{q_-} + q_+ \frac{E\theta_3}{q_-}}$$

where the corresponding integral equations are:

$$p_2(t) = 1 - F_{\theta_{2/q_-}}(t) + \int_0^t p_2(t-s)F\zeta(ds)$$

$$p_3(t) = 1 - F_{q_+\theta_{3/q_-}}(t) + \int_0^t p_3(t-s)F\zeta(ds)$$

So, to solve it practically, we developed approach that supported by elsewhere [6-10]. Using these equations, we can test whether our model describes experimental results from the temperature-sensitive channels TPRV1 and TPRM8. Our model can be used to understand the rate dependence of the dynamics of the channel and to predict the temperature sensitivity of the channel. In order to use our theory, we expressed it into physical model as described below.

The transition between closed state 1 and closed state 2 is defined as:

$$Closed1 \rightarrow Closed2$$

i.e.

$$\psi_0(\tau) = k_c \ \exp(-k_c, \tau) \ and \ \psi_0(\tau) = k_o \ \exp(-k_o, \tau)$$

where $\psi(\tau)$ is the nonexponential distribution and $k_o \ and \ k_c$ is the rate of close or open states. These rates are assumed to depend on an externally applied, time dependent voltage signal. The distributions of residence time intervals implies that the corresponding observed two state dynamics of current fluctuations is not Markovian. This can be expressed as:

$$\psi_0(\tau) = \sum_{i=1}^{N} c_i \, \lambda_i \ \exp(-\lambda_i, t)$$

With weight $c_i$ obeying $\sum_{i=1}^{N} c_i = 1$ . The rationale behind this fitting procedure is the assumption that the corresponding state consists of N discrete sub-states, separated by potential barriers. This method constitutes the working tool for the majority of molecular physiologists in interpreting their experimental data within a discrete Markovian scheme [6-8].

However, the problem with the addition of new states, is that the number of sub-states needed to fit the experimental data can depend on the experimental conditions. For example, the experimental gating dynamics of a Shaker potassium channel has been successfully described by a sequential 8-state Markovian scheme with 7 closed states for a fixed value of temperature about T = 20$^{o}$C [6-8, 11-12]. However, to describe the experimental data over a small extended temperature regime between $10 - 20^{o}$C already necessitates to add three additional closed sub-states [6-8, 11-12]. This set of equations can formally be derived following the approaches described elsewhere [6-10]. So, to develop three states model into three states model we start our channel driving gating with the transition of $Closed1 \leftrightarrow Closed2$ .

$$\psi_{c'}(s) = \frac{k_0^{(0)}(s + k_r)}{(s + k_0^{(0)})(s + k_r) + k_c s}$$

Inversion of this equation yields:

$$\psi_{c'}(\tau) = c_1\lambda_1 \exp(-\lambda_1\tau) + c_2\lambda_2 \exp(-\lambda_2\tau)$$

With the rate coefficient

$$\lambda_{1,2} = \frac{1}{2}(k_0^{(0)} + k_{in} + k_\tau \pm \sqrt{\left(k_0^{(0)} - k_{in} - k_\tau\right)^2 + 4k_0^{(0)}k_{in})}$$

Corresponding weight factors

$$c_{1,2} = \frac{\left[(k_0^{(0)} + k_{in} + k_\tau)\right]}{2\sqrt{\left(k_0^{(0)} - k_{in} - k_\tau\right)^2 + 4k_0^{(0)}k_{in})}}$$

The mean residence time of this equation is:

$$\langle\tau_{c\prime}\rangle = \frac{1}{k_0^{(0)}}(1 + \frac{k_{in}}{k_\tau})$$

And could be characterized as coefficient of variations:

$$C = \frac{\sqrt{\langle\tau^2\rangle - \langle\tau\rangle^2}}{\langle\tau\rangle}$$

Together with the rate coefficient rate equation that appears above, we can write:

$$C_{c'} = \sqrt{1 + \frac{2k_0^{(0)}k_{in}}{(k_{in} + k_\tau)}}$$

In the case of

$$k_\tau \ll k_c \ll k_0^{(0)}$$

We can say that

$$C_{c'} \approx \sqrt{\frac{2k_0^{(0)}k_{in}}{k_{in}}} \gg 1$$

This implies that the distribution has a small but very broad long time tail, which in turn results in long time variance of the residence times. As described by Goychuk et al. [49] [29], the signal to noise ratio is then strongly suppressed in the low frequency limit $\Omega \to 0$ by the factor

$$\frac{1}{N(0)} = \frac{C_0^2 + C_C^2}{2}$$

where $C_0$ is the coefficient of variation for $\psi_0, C_0 = 1$ in the present case. This represents the first clear non-Markovian effect in our model. We also note that the auxiliary functions $\mathbb{N}(s)$ can be recast as

$$\mathbb{N}(s) = \frac{s(s + k_{in} + k_\tau)}{[s + \mu_1(T)][s + \mu_2(T)]}$$

where

$$\mu_{1,2}(T) = \frac{\left[\left(k_0^{(0)} + k_c^{(0)} + k_{in}\right)\right]}{2} + k_\tau \pm \sqrt{k_0^{(0)} + k_c^{(0)} - k_{in} - k_\tau + 4k_0^{(0)}k_{in}}$$

are the decay rate of these of the conductions times correlations. By using the equations of $\mathbb{N}(s)$ together with the first equations here in the process, we obtain:

$$\eta(\Omega, T) = \frac{(\Delta g)^2 q^2 v^2(T)}{16(k_B T)^2 cosh^2\left[\frac{\epsilon(T)}{2k_B T}\frac{\Omega^2 + (k_{in} + k_\tau)^2}{[\Omega^2 + \mu_1^2(T)][\Omega^2 + \mu_2^2(T)]}\right]}$$

and

$$R_{SN}(\Omega,T) = \frac{\pi A^2 q^2 v(T)}{8(k_B T)^2 cosh^2\left[\frac{\epsilon(T)}{2k_B T}\frac{\Omega^2 + (k_{in} + k_\tau)^2}{\Omega^2 + (k_{in} + k_\tau)^2 + k_0^{(0)} k_{in}}\right]}$$

The remaining parameters are:

$$v(\ T) = \langle \pi_0{}^{-1} \rangle + \langle \tau_{c\prime} \rangle^{-1} = k_c^{(0)} + \frac{k_0^{(0)} k_\tau}{(k_{in} + k_\tau)}$$

where

$$k_{c,0}^{(0)} = \ v_0 exp\left[\frac{-\Delta G_{0,c}(v_0)}{(k_B T)}\right]$$

and

$$\epsilon(T) = \Delta G_0(V_0) - \Delta G_C(V_0) + k_B \ln(\frac{1 + k_{in}}{k_\tau})$$

Which leads to exponential Arrhenius rates:

$$k_{in} = v_0 \exp(\frac{-\Delta G_{in}}{k_B \mathrm{T}})$$

and

$$k_\tau = v_0 \exp(\frac{-\Delta G_{in}}{k_B \mathrm{T}})$$

According to these equations we performed numerical calculations for the experimental dataset that we use to calculate $\Delta G, \Delta S\ and\ \Delta H$ in the three states Closed1, Closed2 or Open of TRPV1 and TRPM8 channels.
